# AlphaGenome Enhances Personal Gene Expression Prediction but Retains Key Limitations

**DOI:** 10.1101/2025.08.05.668750

**Authors:** Sana Kashyap, Aarthi Ramakrishnan, Li Shen

**Affiliations:** Bergen County Academies; Department of Artificial Intelligence and Human Health, Icahn School of Medicine at Mount Sinai, New York, USA; Department of Neuroscience, Icahn School of Medicine at Mount Sinai, New York, USA

**Keywords:** gene expression, deep learning, genome AI model, personalized prediction

## Abstract

Recent years have seen rapid growth in sequence-to-function (S2F) models that predict diverse molecular readouts from DNA sequence alone. However, these models have faced criticism for their limited accuracy in predicting individual-level gene expression. AlphaGenome, a state-of-the-art S2F model, excels across a range of sequence-based prediction tasks, but its utility for personal expression prediction remains incompletely assessed. Here, we evaluate AlphaGenomes ability to predict tissue-specific, personal gene expression and find that it significantly outperforms its predecessor, Enformer. Across 1,000 genes and 11,471 eligible genetissue pairs, AlphaGenome significantly outperforms Enformer for 3,103 pairs, whereas Enformer significantly outperforms AlphaGenome for 828 pairs, yielding a win ratio of 3.75. This imbalance is supported by a global permutation test (***P* = 2.0 *×* 10**^***−*4**^), with a median correlation difference of r = 0.046. Nevertheless, pretrained AlphaGenome substantially underperforms Elastic Net and Random Forest trained on individual-level data. These findings demonstrate that AlphaGenome better captures individual-specific gene expression variation than Enformer, while highlighting substantial room for improvement in sequence-based prediction of personal gene expression.

## Introduction

Understanding the relationship between genome sequences and molecular phenotypes– such as chromatin modifications, DNA accessibility, and gene expression–has long been a central focus in genomics. Among these phenotypes, gene expression is arguably the most extensively studied due to its strong associations with health and disease [1]. Over the past few decades, the relationship between genetic variation and gene expression has been studied extensively through expression quantitative trait locus (eQTL) mapping, which typically tests associations between individual genetic variants and expression levels using regression models [2]. Related work has focused on predicting genetically regulated gene expression (GReX) by integrating information across multiple cis-regulatory variants using methods such as penalized linear regression [3–5] and Random Forest [6].

More recently, deep learning-based genome AI models have emerged as powerful alternatives to traditional methods [7–13], demonstrating remarkable performance in predicting a wide range of molecular phenotypes across previously unseen genomic regions. Among these, Enformer [11] represents a landmark advancement, combining convolutional and transformer layers to capture long-range interactions and outperform prior approaches. At the time of its release, Enformer featured significantly more parameters than its predecessors and required substantially greater computational resources for training. Building on this foundation, AlphaGenome [13] is currently the state-of-the-art (SOTA) genome AI model. It employs a 1 Mb context window and delivers predictions at single-basepair resolution, achieving substantial improvements over existing methods across multiple predictive tasks.

AlphaGenome has demonstrated improved performance across several gene-expression-related variant benchmarks, including prediction of eQTL effects, enhancer-gene linking, and polyadenylation QTL effects. For fine-mapped GTEx eQTLs, AlphaGenome increased the tissue-weighted mean Spearman correlation with observed effect sizes from 0.39 to 0.49 and improved the mean auROC for predicting effect direction from 0.75 to 0.80 relative to Borzoi. These results indicate an improved ability to identify the regulatory consequences of individual variants. However, eQTL benchmarks evaluate variants against population-derived association statistics and do not directly establish whether predictions from complete personal genomes can explain measured expression differences across individuals. The present study evaluates this complementary individual-level prediction task.

The impressive performance of genome AI models has sparked growing interest in their potential to predict molecular phenotypes–particularly gene expression–at the individual level. Such capabilities hold significant promise for applications in drug target discovery and precision medicine. However, early investigations have yielded disappointing results: many of Enformer’s predictions were negatively correlated with observed expression levels [14, 15]. In response, researchers have attempted to fine-tune Enformer and similar models using individual-specific data, yielding modest but encouraging improvements [16, 17].

This study evaluates the potential of AlphaGenome for predicting personal gene expression in a zero-shot setting, without training or fine-tuning the model on paired individual-level genome and expression data. Although AlphaGenome was not explicitly trained on individual-level data, its enhanced architecture and expanded training set motivate testing whether its out-of-the-box predictions improve upon those of its predecessor. To assess its capabilities and limitations, we compare AlphaGenome with Enformer as well as two conventional machine-learning methods for GReX prediction: Elastic Net and Random Forest. In addition to the main evaluation, we examine exploratory subsets of genetissue pairs in which Random Forest and Elastic Net show discordant performance and conduct case studies to examine the relationship between genetic variants and AlphaGenomes individual-level predictions.

## Results

### AlphaGenome Significantly Outperforms Its Predecessor in Predicting Personal Gene Expression

The GTEx database [18], which includes RNA-seq data from 953 individuals across 50 tissues, was used as a benchmark to evaluate each models ability to predict personal gene expression. To account for known technical and biological confounders, normalized gene expression values were adjusted using GTEx-provided covariates, including genotype principal components, inferred expression covariates, sex where applicable, PCR amplification, and genotyping platform. The resulting residualized expression values were used as the observed outcomes for all four models, ensuring a consistent evaluation target. The regressed expression matrix, downloaded from the GTEx website, contains 42,372 genes. After filtering for sparsity, 377,857 gene-tissue pairs remained for model training and evaluation. Both Elastic Net and Random Forest were trained and evaluated separately for each gene using nested 10-fold cross-validation. Based on the results from Elastic Net, 1,000 genes were randomly selected to span a range of coefficients of determination (R2) across tissuesfrom negative values to greater than 0.2.

AlphaGenome was originally trained to predict population-averaged expression levels for all GTEx tissues; thus, its predictions for individual sequences on the selected 1,000 genes were used directly. Since Enformer does not natively output GTEx tissue predictions, we followed AlphaGenome’s approach: Enformer’s frozen embeddings were used to train predictors for population-averaged expression across all GTEx tissues, and these predictors were then applied to individual sequences. For fair comparison across models, Pearson correlation was used as the evaluation metric, as it does not require scale-matching between predicted and observed values. For details of data preprocessing, model training and evaluation, see Section S1.

Figure 1 presents the distribution of Pearson correlation coefficients across all 11,471 genetissue pairs for the four methods compared. As expected, Elastic Net and Random Forest achieved higher median correlations (0.166 and 0.144, respectively) than AlphaGenome (0.041) and Enformer (0.006), reflecting the advantage of models trained directly on individual-level expression data. Nevertheless, AlphaGenome consistently exhibited a more favorable distribution of correlations than Enformer, with a higher median correlation and a more pronounced positive tail. AlphaGenome yielded 2,459 significantly positive and 971 significantly negative correlations, compared with 1,557 significantly positive and 1,873 significantly negative correlations for Enformer. These findings suggest that, despite its limited overall predictive accuracy, AlphaGenome captures individual-level expression variation more effectively than its predecessor without fine-tuning on paired personal genomes and transcriptomes.

**Fig. 1.**
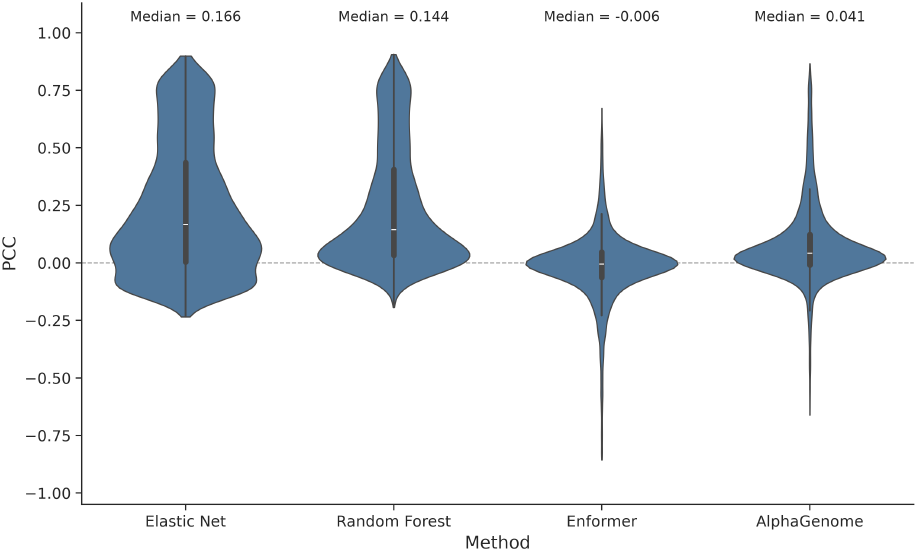
Violin plots of Pearson correlations of the four methods across all gene-tissue pairs.

To further assess their relative performance, head-to-head comparisons were conducted between AlphaGenome and each of the other three methods across all gene-tissue pairs. AlphaGenome significantly outperforms Elastic Net in 1,048 gene-tissue pairs (Figure 2b) and Random Forest in 221 gene-tissue pairs (Figure 2c). When compared to Enformer, AlphaGenome shows a clear advantage: 3,103 gene-tissue pairs have significantly higher correlations than Enformer, while only 828 favor Enformer, resulting in a winning ratio of 3.75 (Figure 2a).

**Fig. 2.**
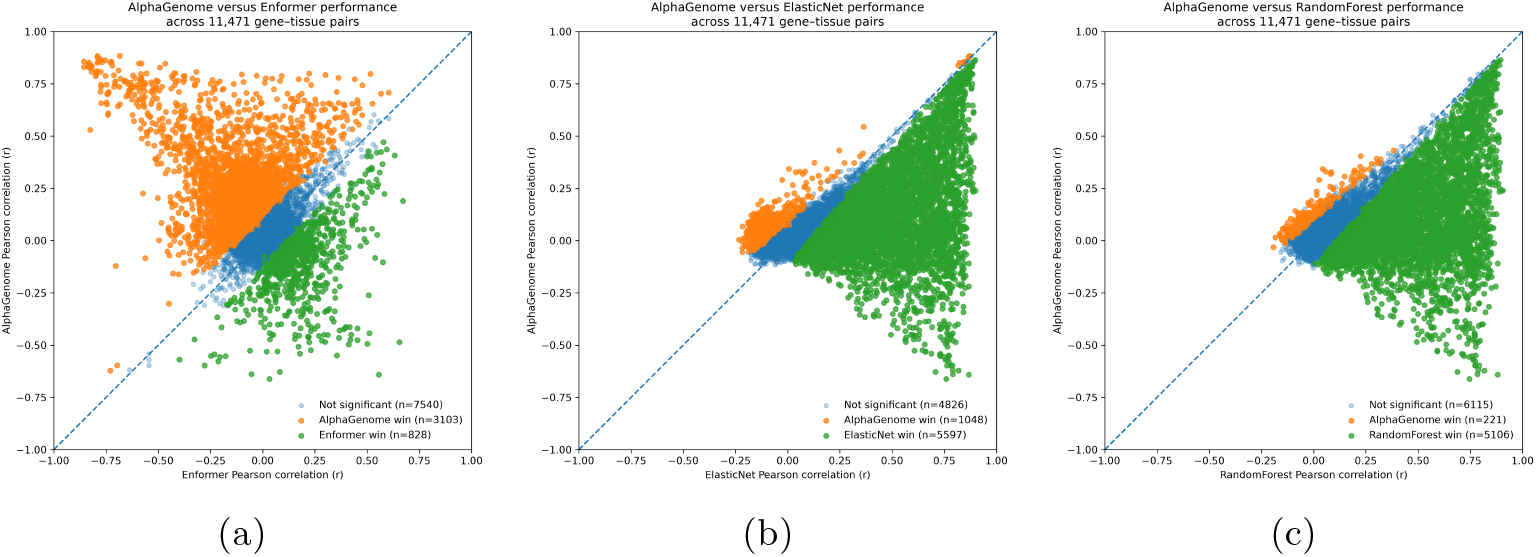
AlphaGenome compared with Enformer, Elastic Net, and Random Forest for personal gene expression prediction. Each point is one of 11,471 gene-tissue pairs, plotted as the Pearson correlation between predicted and observed expression for AlphaGenome (y-axis) against the comparison method (x-axis); the dashed line marks equal performance. Points above the diagonal favor AlphaGenome. Pairs are colored by: orange, AlphaGenome significantly better; green, the comparison method significantly better; blue, not significant. **(a)** AlphaGenome vs. Enformer: AlphaGenome wins 3,103 pairs and Enformer 828. (b) AlphaGenome vs. Elastic Net **(c)** AlphaGenome vs. Random Forest Elastic Net and Random Forest were trained directly on individual-level data, whereas AlphaGenome was not.

To investigate whether AlphaGenome’s performance advantage over Enformer was consistent across biological contexts, we compared the mean paired differences in Pearson correlation between AlphaGenome and each benchmark model across 15 GTEx tissues (Figure 3). AlphaGenome consistently outperformed Enformer across all evaluated tissues, with positive mean differences in Pearson correlation and 95% bootstrap confidence intervals excluding zero. The magnitude of improvement varied by tissue, with the largest advantage observed in cultured fibroblasts. In contrast, AlphaGenome underperformed both Elastic Net and Random Forest across all 15 tissues, with negative mean paired differences throughout. These results demonstrate that AlphaGenome’s improvement over Enformer is broadly consistent across diverse tissue types, although supervised models trained on individual-level expression data continue to achieve superior predictive performance.

**Fig. 3.**
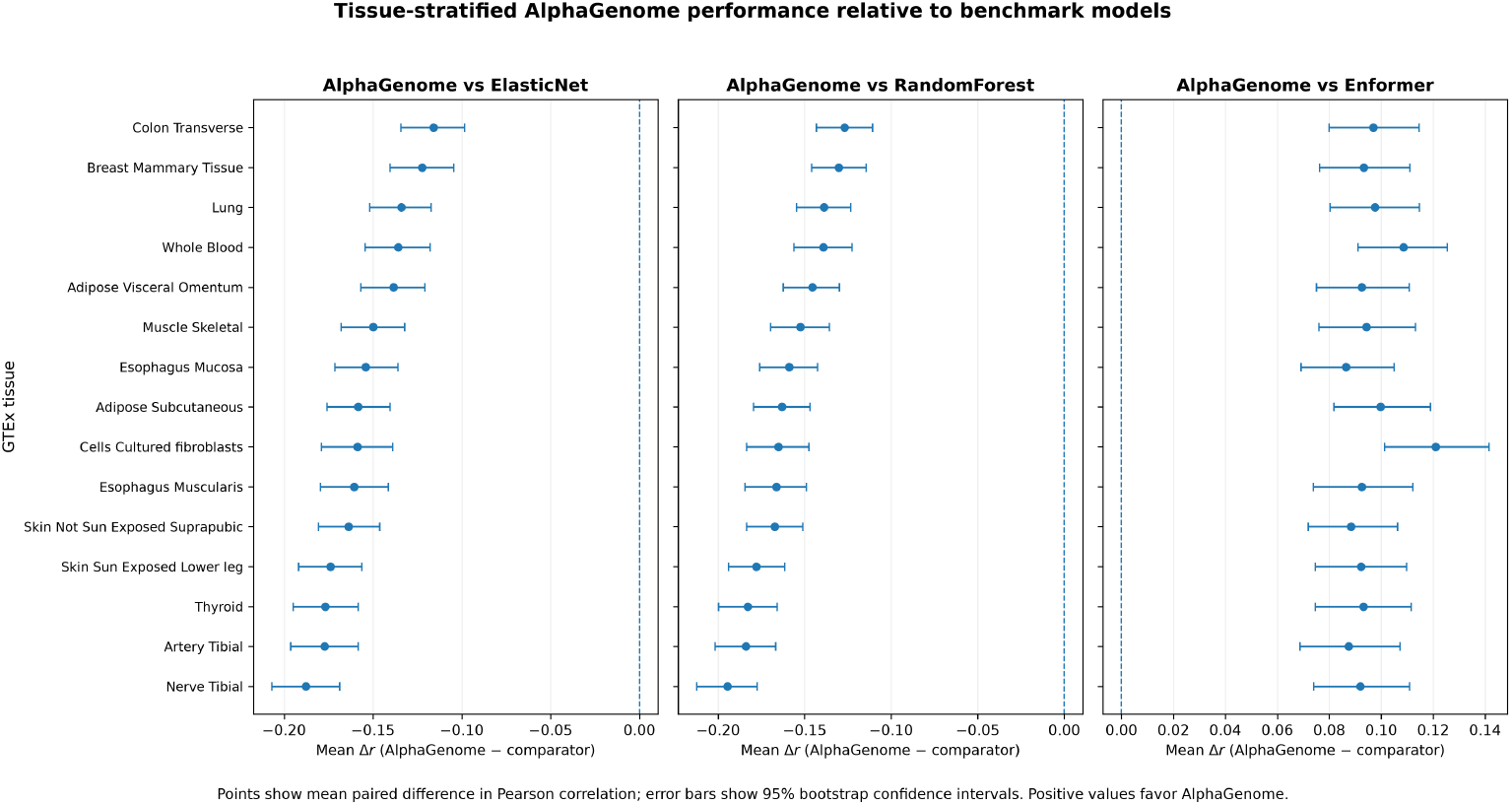
Tissue-specific performance of AlphaGenome relative to comparison models. Each point shows the mean paired difference in Pearson correlation (AlphaGenome minus comparison model) for one tissue, and horizontal error bars show 95% bootstrap confidence intervals. The dashed vertical line at zero indicates equal performance. Positive values favor AlphaGenome and negative values favor the comparison model.

However, while AlphaGenome retained significant advantages over all three comparator models, this pattern depended strongly on baseline gene predictability (Figure 4) Relative to ElasticNet, 1,048 gene–tissue pairs significantly favored AlphaGenome, with 89.3% of these wins occurring in the two lowest ElasticNet predictability strata (*R*^2^ *≤* 0.01). Similarly, 221 gene–tissue pairs significantly favored AlphaGenome over Random Forest, of which 86.9% occurred among genes with baseline *R*^2^ *≤* 0.05. In contrast, AlphaGenome showed 3,103 significant wins over Enformer, and these were concentrated among more predictable genes: 76.0% occurred in the *R*^2^ *>* 0.05 strata.

**Fig. 4.**
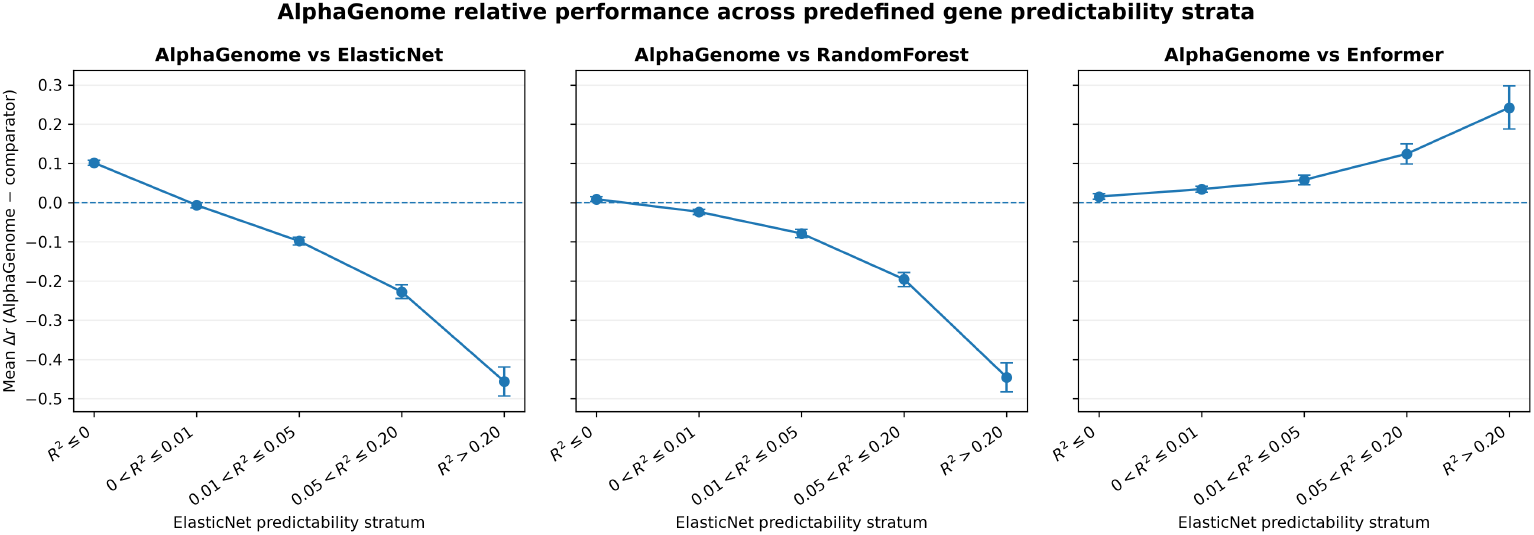
AlphaGenome performance across gene predictability strata. Genetissue pairs were stratified into five predictability categories based on Elastic Net cross-validated (): (*R*^2^*≤* 0), (0 ¡ *R*^2^ *≤* 0.01), (0.01 *< R*^2^ *≤* 0.05), (0.05 *< R*^2^ *≤* 0.20), and (*R*^2^ *>* 0.20). Points represent the mean paired difference in Pearson correlation ((Δ*r*)) between AlphaGenome and Elastic Net (left), Random Forest (middle), or Enformer (right), with error bars indicating 95% confidence intervals. Positive values indicate superior performance by AlphaGenome, whereas negative values favor the comparator.

To further characterize differences in predictive performance between AlphaGenome and Enformer, we stratified genetissue pairs according to the direction and statistical significance of each model’s Pearson correlation with observed expression (Figure 5). Among the 11,471 evaluated genetissue pairs, 1,471 exhibited significantly positive correlations for AlphaGenome but significantly negative correlations for Enformer, compared with only 264 pairs showing the reverse pattern. Of the former, AlphaGenome significantly outperformed Enformer in 1,470 pairs, whereas Enformer significantly outperformed AlphaGenome in all 264 pairs exhibiting the opposite correlation pattern. This pronounced asymmetry suggests that AlphaGenome more frequently recovers positive associations with observed expression in cases where Enformer produces significant negative correlations. Nevertheless, 5,080 pairs exhibited nonsignificant correlations for both models, highlighting the limitations of both approaches in predicting individual-level expression variation. Together, these findings demonstrate that AlphaGenome’s improvements over Enformer are not uniformly distributed across genes but are particularly pronounced in cases involving reversals in correlation direction.

**Fig. 5.**
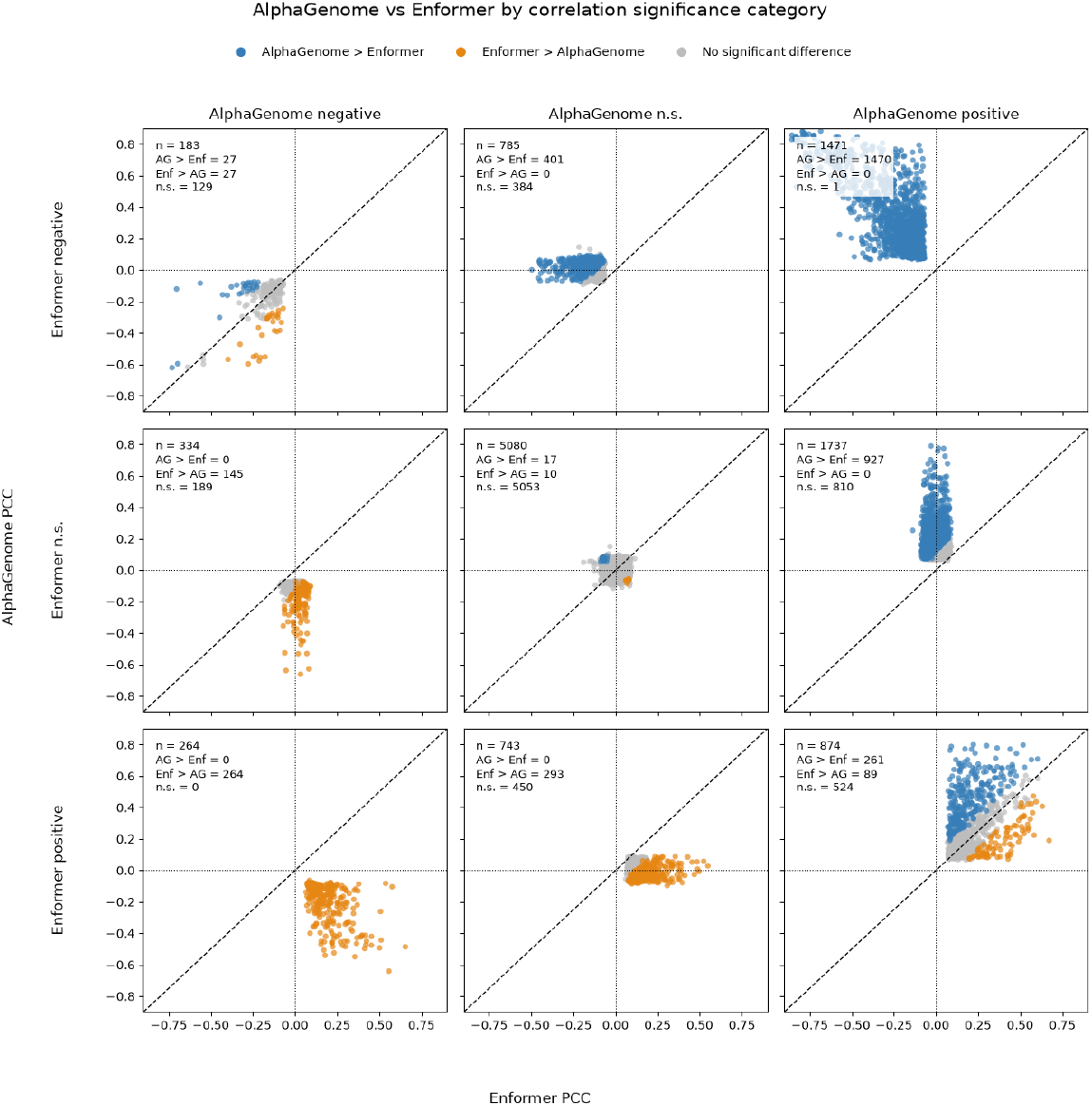
Comparison of AlphaGenome and Enformer predictive performance across correlation significance categories. Genetissue pairs were classified into nine categories based on the direction and statistical significance of Pearson correlations between predicted and observed expression for AlphaGenome and Enformer. Correlations were categorized as significantly positive, significantly negative, or nonsignificant. Each panel represents one combination of Enformer (rows) and AlphaGenome (columns) correlation categories, with the number of genetissue pairs indicated. Within each panel, points are colored according to the outcome of pairwise permutation testing with BenjaminiHochberg correction, distinguishing pairs significantly favoring AlphaGenome, significantly favoring Enformer, or exhibiting no significant difference.

### AlphaGenome Captures Genotype-Associated Expression Patterns with Limited Residual Predictive Power

To investigate the contribution of genotype-associated variation to AlphaGenome’s predictive performance, we examined 20 selected genetissue pairs in which AlphaGenome significantly outperformed Enformer and achieved a Pearson correlation of at least 0.30. After adjusting both predicted and observed expression for lead-variant genotype dosage, squared Pearson correlations decreased across all 20 pairs (Figure 6). The median squared correlation declined from 0.570 before adjustment to 0.005 afterward, corresponding to a median pairwise relative reduction of 99.1%. These findings suggest that AlphaGenome’s predictive associations in these selected examples were predominantly attributable to differences associated with the lead genotype, with substantially weaker associations remaining after genotype adjustment. However, because the analysis focused on selected high-performing genetissue pairs, the findings do not necessarily generalize to the full evaluation dataset.

**Fig. 6.**
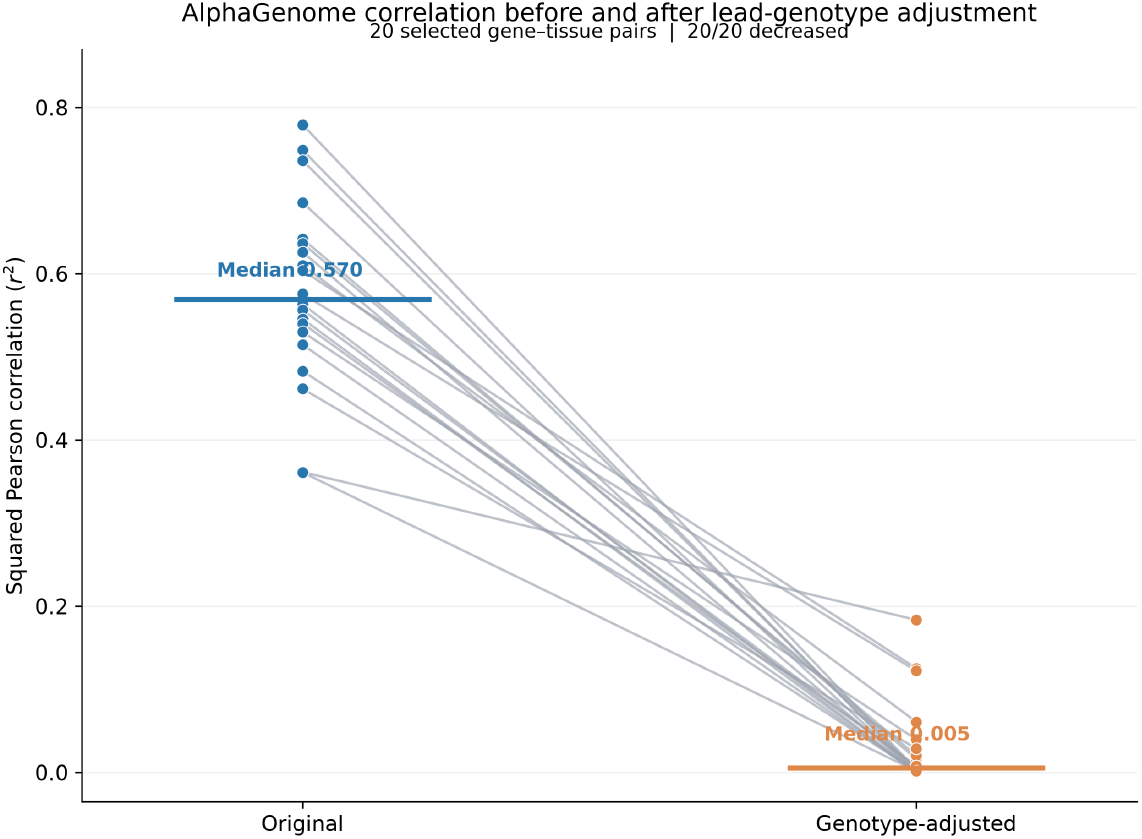
Effect of genotype adjustment on AlphaGenome’s predictive performance across 20 selected genetissue pairs. Paired comparison of squared Pearson correlations ((*R*^2^)) between AlphaGenome-predicted and observed expression before and after adjustment for lead-variant genotype dosage. Each connected pair of points represents one genetissue pair, with horizontal lines indicating median (*R*^2^).

## Discussion

### Is bigger always better?

For researchers in the field of large language models (LLMs), the answer is often a resounding yes, as the scaling law has consistently held true [18]. We are currently witnessing the rise of extreme-scale LLMs–such as GPT-4 [19] and Grok 4 [20]–that continue to grow in size and capability. A similar trend may be unfolding in genomics, as the declining cost of sequencing paves the way for personal genomes to be generated at unprecedented scale [21].

AlphaGenome, with its 1 Mb context window and single-basepair resolution, represents the current state-of-the-art in genome AI. It has demonstrated exceptional accuracy in predicting molecular phenotypes across multiple modalities using the reference genome. As shown in this study, AlphaGenome also improves individual-level gene expression prediction relative to Enformer, although the overall improvement in correlation is modest and its performance remains below that of models trained directly on individual-level data. This finding is consistent with AlphaGenomes previously reported improvements in predicting eQTL effect magnitude and direction. Nevertheless, accurate prediction of individual variant effects does not necessarily translate into accurate prediction of expression across individuals, which requires integrating the combined effects of multiple variants within each personal genome. Although AlphaGenomes architectural advances, expanded context window, and multimodal training may contribute to its improved performance, the present study does not isolate the contributions of these individual features.

Despite these advances, AlphaGenome still performs less favorably than conventional machine-learning models trained directly on individual-level data. Recent studies have begun to address this limitation by training or fine-tuning deep sequence models on paired personal genomes and transcriptomes. Variformer and fine-tuned Enformer improve prediction for held-out individuals at genes represented during training, while p-SAGE-net provides a scalable framework for personal-genome training. However, these studies also report limited generalization to genes or loci not represented during training. [17, 22, 23] These supervised approaches address a different question than the present study, which evaluates AlphaGenomes transfer to personal expression prediction without individual-level fine-tuning. Fine-tuning AlphaGenome using paired personal genomes and transcriptomes may further improve its predictive performance and represents an important direction for future investigation.

This gap reflects, in part, a key limitation of the study: the model-training setups differ in training data and task. AlphaGenome and Enformer are pretrained sequence-to-function models whereas the ElasticNet and Random Forest models were trained for the expression-prediction task. The comparisons are informative about whether AlphaGenome performs better where nonlinear modeling helps, but they are not controlled head-to-head comparisons.

Another limitation of the present study is that model evaluation was restricted to 1,000 selected genes, which does not constitute a genome-wide benchmark. To improve representation across the full spectrum of baseline gene predictability, the gene set was constructed using stratified sampling based on out-of-fold Elastic Net performance. Nevertheless, the conclusions are limited to genes satisfying the studys expression and missingness criteria. Model performance also varied substantially across baseline predictability strata and tissues, highlighting the importance of evaluating predictive accuracy across different genetic contexts rather than relying exclusively on aggregate results.

Finally, the current genotype dosage analysis is selective and limited in scope. While the top 20 gene-tissue pairs where AlphaGenome significantly outperformed Enformer were selected, these cannot characterize every pair. However, these cases are useful for understanding a potential mechanism behind high predictive correlations. Future research should expand this analysis systematically across many more genes and ask how much predictive signal remains after major cis-regulatory effects are removed.

## Acknowledgements

This work was supported in part through the Minerva computational and data resources and staff expertise provided by Scientific Computing and Data at the Icahn School of Medicine at Mount Sinai and supported by the Clinical and Translational Science Awards (CTSA) grant UL1TR004419 from the National Center for Advancing Translational Sciences.

## S1 Supplementary Methods

### Data Preprocessing

For RNA-seq data, GTEx v10 tissue-specific normalized gene expression phenotypes and corresponding covariate matrices were obtained from the GTEx portal. Within each tissue, individual identifiers were aligned between the expression and covariate matrices. For each gene, normalized expression was regressed on the GTEx-provided covariates, including five genotype principal components, a tissue-specific number of inferred expression covariates, sex where applicable, and indicators for PCR amplification and genotyping platform. The resulting residuals, representing expression variation after adjustment for these covariates, were used as the observed expression outcomes throughout the study, including gene selection and evaluation of AlphaGenome, Enformer, Elastic Net, and Random Forest. Genetissue pairs with more than 50% missing expression values were excluded.

For personal genomes, phased whole-genome sequencing data (V9) were used. The genotype matrices used for Elastic Net and Random Forest were constructed as follows. First, for each gene, variants within a 1 Mb window centered on the transcriptional start site (TSS) were extracted from phased variant call format (VCF) files separately for each haplotype. Second, each variant was encoded in binary form (0/1) per haplotype and concatenated to generate individual-level numerical feature vectors for downstream model training. Both SNVs and indels were included, and variants with a minor allele frequency below 1% were filtered out. For AlphaGenome and Enformer, the haplotype sequences were generated using the *consensus* command from the bcftools[24] package. Both haplotypes were used as input to the deep learning models, and the resulting outputs were averaged to produce the final predictions.

### Use AlphaGenome and Enformer for Personal Expression Prediction

For AlphaGenome, a 1 Mb window centered on the transcriptional start site (TSS) of each gene was used as input. The model outputs predictions over the same window at single-basepair resolution. For each gene, predicted expression values from the TSS to the transcriptional end site (TES) were averaged to represent that gene’s expression level. AlphaGenome’s half-window size of 500 Kb is sufficient to span the full gene body (TSS to TES) for the vast majority of genes; only 422 genes–approximately 0.7% of all GTEx genes–exceed 500 Kb in length.

Enformer, by contrast, requires training to predict GTEx tissue-level expression. The same train-validation-test split as used in the Borzoi [12] model was adopted, where each GTEx gene was assigned to a set based on the genomic region closest to its TSS. A 196,608 bp window centered on the TSS was extracted from the reference genome and used as input to the pretrained Enformer to obtain an embedding. As with AlphaGenome, predicted values from TSS to TES were averaged to represent gene-level expression. The target values were defined as the average expression across all GTEx individuals for each tissue.

The position-averaged embedding vectors were then used to train a Ridge regressor to predict tissue-specific expression levels. The regularization strength was selected from 30 evenly spaced values in log space between 1 and 5. The optimal model was chosen based on R2 performance on the validation set. For personal-level expression prediction, individual genome sequences were fed into Enformer to generate embeddings, which were then input into the trained Ridge regressor to produce tissue-specific expression predictions.

### Use Elastic Net and Random Forest for Personal Expression Prediction

The genotype matrices were used as input to Elastic Net and Random Forest. Model performance was evaluated using 10-fold nested cross-validation. Within the inner loop, Elastic Net hyperparameters were optimized using 5-fold cross-validation to select the regularization parameter from 30 candidate *α* values. Random Forest models were trained using default hyperparameter settings. All analyses were conducted using the scikit-learn package [25].

### Choosing 1,000 Genes that Span Different R2 Values

Five total strata were created based on out-of-fold Elastic Net’s performance on each gene: *R*^2^ *<* 0, 0 *< R*^2^ ≤ 0.01, 0.01 *< R*^2^ *≤* 0.05, 0.05 *< R*^2^ *≤* 0.20, *R*^2^ *>* 0.20. 200 genes were randomly selected from each strata to create a set of 1,000 genes (Supplemental Table 1) that span a wide range of predictability levels

### *In silico* mutagenesis

*In silico* mutagenesis (ISM) was performed to estimate the marginal effect of individual genetic mutations on AlphaGenome’s output. Each mutation was introduced into a reference sequence spanning 1 Mb centered on the TSS, and the resulting predicted expression was compared to the prediction from the unmodified reference sequence. The difference between these two predictions was taken as the mutation’s effect.

For Random Forest, the reference expression was obtained using a feature vector of all zeros. Each mutation was then individually activated (set to one) to compute the corresponding alternative expression. To enable direct comparison between AlphaGenome and Random Forest, the marginal effects for all mutations were rescaled to the [0, 1] range.

### Paired Permutation Test

For each eligible gene–tissue pair *k* = 1, …, *K*, we computed the difference in predictive correlation between Methods A and B on the same individuals:

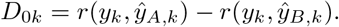

Positive values favor A and negative values favor B. Eligibility criteria and the FDR level were fixed before testing.

#### Permutation scheme

For each permutation, we swapped the complete A and B prediction profiles for a random half of individuals (a fair coin per individual), using the same swap across all genes and tissues to preserve dependence among outputs. All *K* differences were then recomputed. We generated two independent sets of permutations: a calibration bank (*B*_cal_ permutations) and an evaluation bank (*B*_eval_ permutations).

#### Pair-level tests

The calibration bank gives a null distribution 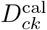 for each pair. Two-sided p-values were

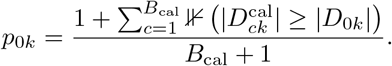

We applied Benjamini–Hochberg correction across all *K* pairs (*q* = 0.05) and labeled significant pairs as A wins (*D*_0*k*_ *>* 0) or B wins (*D*_0*k*_ *<* 0), giving counts *N*_*A*,0_ and *N*_*B*,0_.

#### Global test

For each evaluation permutation *b*, we compared its differences 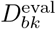 against the same fixed calibration distributions to obtain p-values, applied BH, and counted *N*_*A,b*_ and *N*_*B,b*_. This repeats the full analysis under the null without nested permutations. The test statistic was *T* = *N*_*A*_ *− N*_*B*_, with

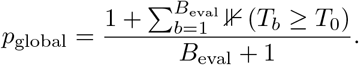

The ratio *N*_*A*,0_*/N*_*B*,0_ is reported descriptively only.

#### Cost and limitations

This design needs 1 + *B*_cal_ + *B*_eval_ correlation recalculations rather than *B*_cal_*B*_eval_. It assumes A and B predictions are exchangeable within individuals under the null. Non-significant pairs are not claimed to be equal.

